# Sex differences in olfactory behavior and neurophysiology in Long Evans Rats

**DOI:** 10.1101/2024.05.21.595192

**Authors:** Kruthika V Maheshwar, Abigail E. Stuart, Leslie M. Kay

## Abstract

In many species, olfactory abilities in females are more sensitive than those in males. Studies in humans show that women have lower olfactory thresholds and are better able to discriminate and identify odors than men. In mice, odorants elicit faster activation from a larger number of olfactory sensory neurons in females than males. Our study explores sex differences in olfaction in Long Evans rats from a behavioral and electrophysiological perspective. Local field potentials (LFPs) in the olfactory bulb (OB) represent the coordinated activity of bulbar neurons. Olfactory gamma (65-120 Hz) and beta (15-30 Hz) oscillations have been functionally linked to odor perception. Spontaneous and odor-evoked OB LFPs were recorded from awake rats at the same time for 12 days. Odors used included urine of both sexes and monomolecular odorants characterized previously for correlation of volatility with behavior and OB oscillations. Sampling duration, baseline gamma and beta power, and odor-elicited beta and gamma power were analyzed. We find that females sample odorants for a shorter duration than males (close to 1s difference). While baseline gamma and beta power do not show significant differences between the two sexes, odor-elicited gamma and beta power in females is significantly lower than in males. Neither sampling duration nor beta and gamma power in females varied systematically with day of estrus. We further verify that variance of these behavioral and physiological measures is not different across sexes, adding to growing evidence that researchers need not be concerned about often-claimed additional variance in female subjects.

**New and Noteworthy:** Olfaction plays a large role in evolutionary processes. However, we know little about sex differences in olfactory bulb neurophysiology, and many scientists believe that females are more variable due to estrus. We show that female rats sniff odors for shorter durations than males and have lower power in neural oscillations related to cognition. Estrus was not related to variance in any measures. Finally, males and females show equal variance on these behavioral and physiological processes.

## Introduction

Females across several species show greater sensitivity or better performance on some measures of olfactory behavior than males. According to a 2019 meta-analysis study in humans, where olfactory discrimination and identification and odor threshold (the lowest perceptible odorant concentration) were examined, women outperformed men in all three aspects (Sorokowski et al., 2019). There may be evolutionary origins for olfactory sex differences. In female pigs, the odor threshold for specific pheromone detection is five times lower than that in males, and this observation has been linked to sexual selection theory (Dorries et al., 1995; Trivers, 1972). Female energetic investment is typically higher in terms of gamete production, gestation (in species where it is applicable), and parental care. More sensitive or precise olfactory ability in females might therefore aid mate selection. Indeed, female rats prefer the urine of healthy males over that of males infected with parasites (Willis & Poulin, 2000).

Behavioral sex differences can often be linked to underlying physiological differences. Studies dealing with mechanistic explanations of sex differences in olfaction have explored cellular, hormonal, and anatomical aspects. Odorants elicit faster responses from a greater number of olfactory sensory neurons in female mice relative to males. Interestingly, in the absence of circulating hormones, the above effect is reversed between the sexes (Kass et al., 2017). The human olfactory bulb (OB) is sexually dimorphic, with females having a larger number of neurons and glia than males (Oliveira-Pinto et al., 2014). Given the above sex differences, it is likely that these may be accompanied by differences in the physiology of the central olfactory system. We tested this hypothesis from the perspective of neuronal oscillations, which are a signature of sensory and perceptual processing in the mammalian OB (Frederick et al., 2016; Kay, 2014; Kay et al., 2009; Rojas-Líbano & Kay, 2008). The roles of synapse and input strength in these oscillations is well-known (Bathellier et al., 2006; Li & Cleland, 2017; Osinski & Kay, 2016; Rosero & Aylwin, 2011). To our knowledge, sex differences in OB oscillations have not been reported.

Neuronal oscillations are rhythmic, repetitive, and coordinated electrical activity of neurons that constrain spike timing in large populations of interacting neurons. This study focuses on beta (15-30 Hz) and gamma (65-120 Hz) oscillations of the OB local field potential (LFP). Beta oscillations are produced in response to repeated presentations of highly volatile odorants such as toluene and hexanone (Lowry & Kay, 2007), are associated with odor-directed motor behavior (Hermer-Vazquez et al., 2007), and increase with learning in operant odor discrimination tasks involving reward-based learning (Frederick et al., 2016; Martin et al., 2004, 2007). OB gamma oscillations are associated with olfactory discrimination in rats and mice (Beshel et al., 2007; Frederick et al., 2016; Lowry & Kay, 2007; Nunez-Parra et al., 2013), as are analogous oscillations in insects (Kay, 2015; Kay & Stopfer, 2006; Patel & Rangan, 2021).

The present study tests for sex differences in 1) odor sampling behavior and 2) OB neural oscillation power, specifically beta and gamma oscillations, at baseline and during odor sampling. OB LFPs from awake, unrestrained rats were recorded during passive odor presentation. A total of seven odorants, five monomolecular and male and female urine, were used. The monomolecular odors span a range of volatilities, evoking neural oscillations and behavior in male rats that have been characterized in previous work (Lowry & Kay, 2007). Male and female rat urine are ethologically relevant cues (Brown, 1991; Osada et al., 2009; Tsai et al., 1994; Zhang & Zhang, 2011) and were included because urine-mediated olfactory communication has the potential to exhibit sex differences. Responses to blank swabs (swabs with no odorants) were also recorded. We found a sex difference in odor investigation time, with females sampling odorants for shorter durations than males early in series of odor presentations. There were no significant differences between sexes in baseline gamma or beta power, but odor-elicited gamma and beta power was lower in females than males. We found no evidence that either sampling duration or oscillations varied relative to estrus day in females differently from males.

## Methods

### Subjects

Adult Long Evans male and female rats (8-10 weeks old at the time of electrode implant; purchased from Envigo), *n*=10 (5F), were used for the experiments. Rats with electrode implants were housed singly on a 14:10 h light/dark schedule (lights on at 7AM and off at 9PM CST) and had unlimited access to food and water. All procedures conformed to Association for Assessment and Accreditation of Laboratory Animal Care (AAALAC) guidelines and were done with approval by The University of Chicago Institutional Animal Care and Use Committee under veterinary supervision.

### Electrode Implants

Electrode implant surgeries were performed under anesthesia. The rats were given a presurgical subcutaneous dose of ketamine cocktail (35 mg/kg ketamine, 5 mg/kg xylazine, and 0.75 mg/kg acepromazine), after which the anesthetized state was maintained by intraperitoneal dosing with ketamine (35 mg/kg) every 45 min to one hour, as determined by changes in respiratory rate. The rats were surgically implanted with bipolar stainless-steel electrodes bilaterally in the OBs (8.5 mm anterior to bregma, 1.5 mm lateral, ∼4 mm deep), straddling the ventral mitral cell layer. Ground and reference screws with wires attached were placed in the right and left posterior dorsal skull. Electrode pins were inserted into a round plastic head stage connector (Ginder Scientific), which was then embedded and secured to the rat’s skull with dental cement. Post-surgically, the rat was monitored for surgical wound resolution, alertness, and food and water intake. The recovery period typically spanned two weeks to rule out any possible sex differences in recovery after which rats were used for the experiment.

### Recording and Odor Presentation

For each experimental session, the rat was placed in a clean polycarbonate cage, 430 x 290 x 201mm (L X B X H), with the head stage cable (NB Labs, 9 pin custom made cable) connected to the recording setup and were allowed to move freely. Each session was designed to contain 7 blocks (corresponding to the 7 odorants; Table 1). Each block consisted of eight trials of a single pure odorant on a cotton swab followed by a blank swab, followed by a 1-minute wait period. The intertrial interval within blocks was 20 s. All rats were recorded for 12 sessions, one session per day, between 10:00 and 13:00 CST, during the rat’s light phase. The order of presentation of odors was randomized across sessions and rats, with the exception that urine odors were presented in succession (Fig. 1). Odors were presented by holding an odorant saturated cotton swab under the nose of the rat. Due to a miscommunication, two slightly different presentation paradigms were used. For the first paradigm, n=6 rats (3F), the first trial always lasted for 10 seconds; for the next 7 trials the cotton swab was held under the snout for 10 s or until the rat rejected the swab, whichever was first (typically, rats do not sample longer than 10s on presentations after the first trial, and sampling drops considerably after trial 2 (Lowry & Kay, 2007). Rejection of the odor was defined as backing away or turning the head away from the swab. In the second paradigm, for n=4 rats (2F), the rats were allowed to sample freely until they rejected the swab on all trials. The beginning and end of odor presentation was recorded by the experimenter by tapping on a piezoelectric strip. LFP data and the piezo signal were recorded using Cheetah 5.0 (Legacy) 5.7.4 - Neuralynx5 at a sampling rate of 2020 Hz with analog filters set at 1-475 Hz.

**Figure 1:**
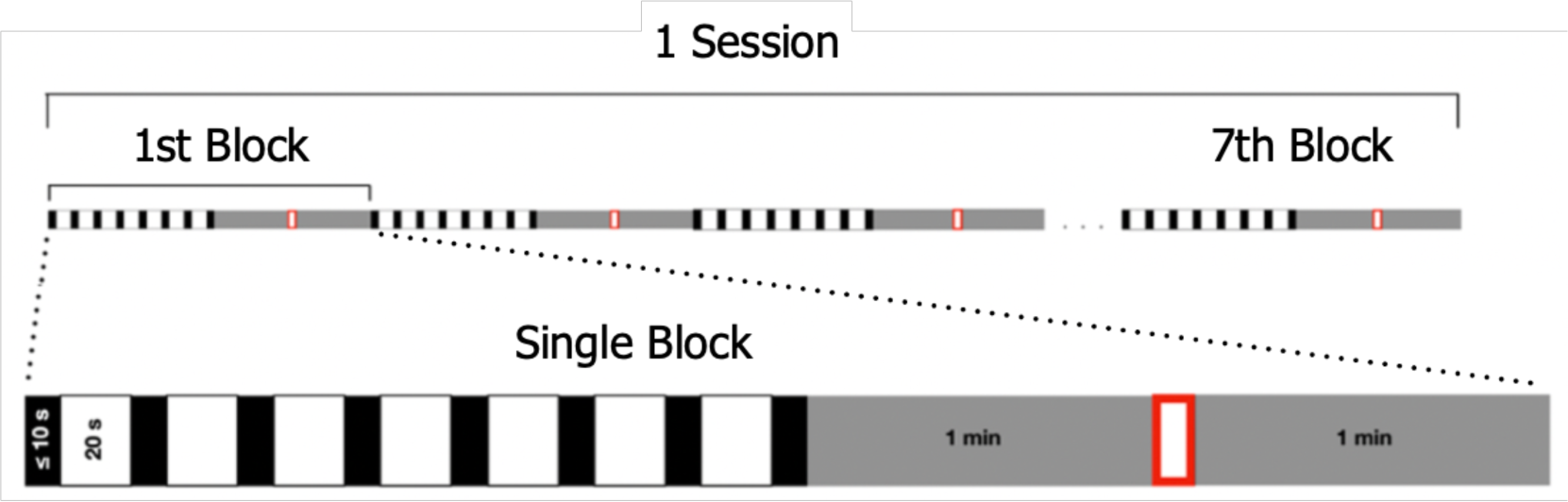
Schematic Representation of a Single Session: Black boxes represent odor presentation trials, white boxes represent the intertrial intervals, and the red box represents the blank swab presentation. Gray boxes represent intervals flanking the blank swab trial.

**Table 1.**
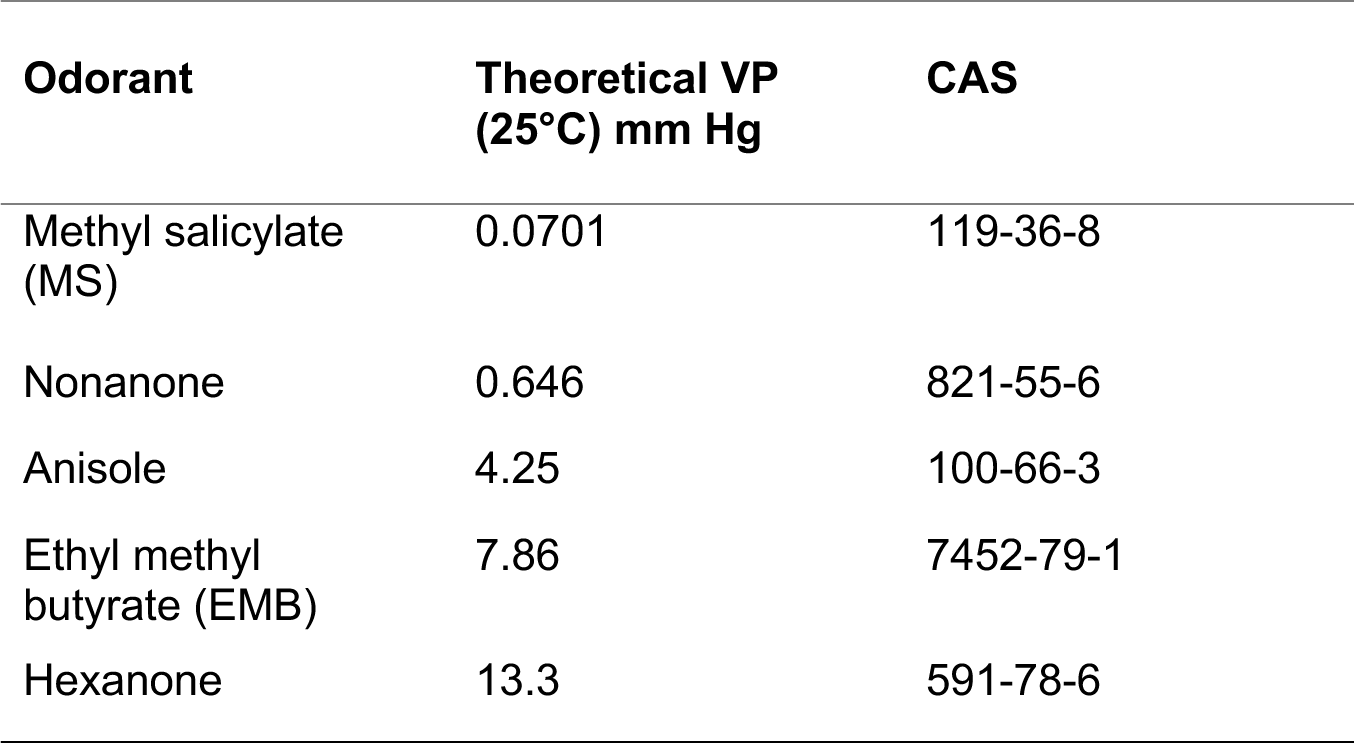
Odors and volatility.

### Estrus Staging

A protocol previously described by (Ajayi & Akhigbe, 2020) to stage mice in the estrus cycle was used to determine the estrus stages of the female rats in the experiment. At the end of each session, female rats were gently restrained by holding them down with the non-dominant hand, while lifting their tails to obtain digital images of the vulva. These images were later used to determine estrus stages, based on the visual features described in Table 2. To ensure experimental consistency, the same maneuver was also carried out on the male rats in the study.

**Table 2.**
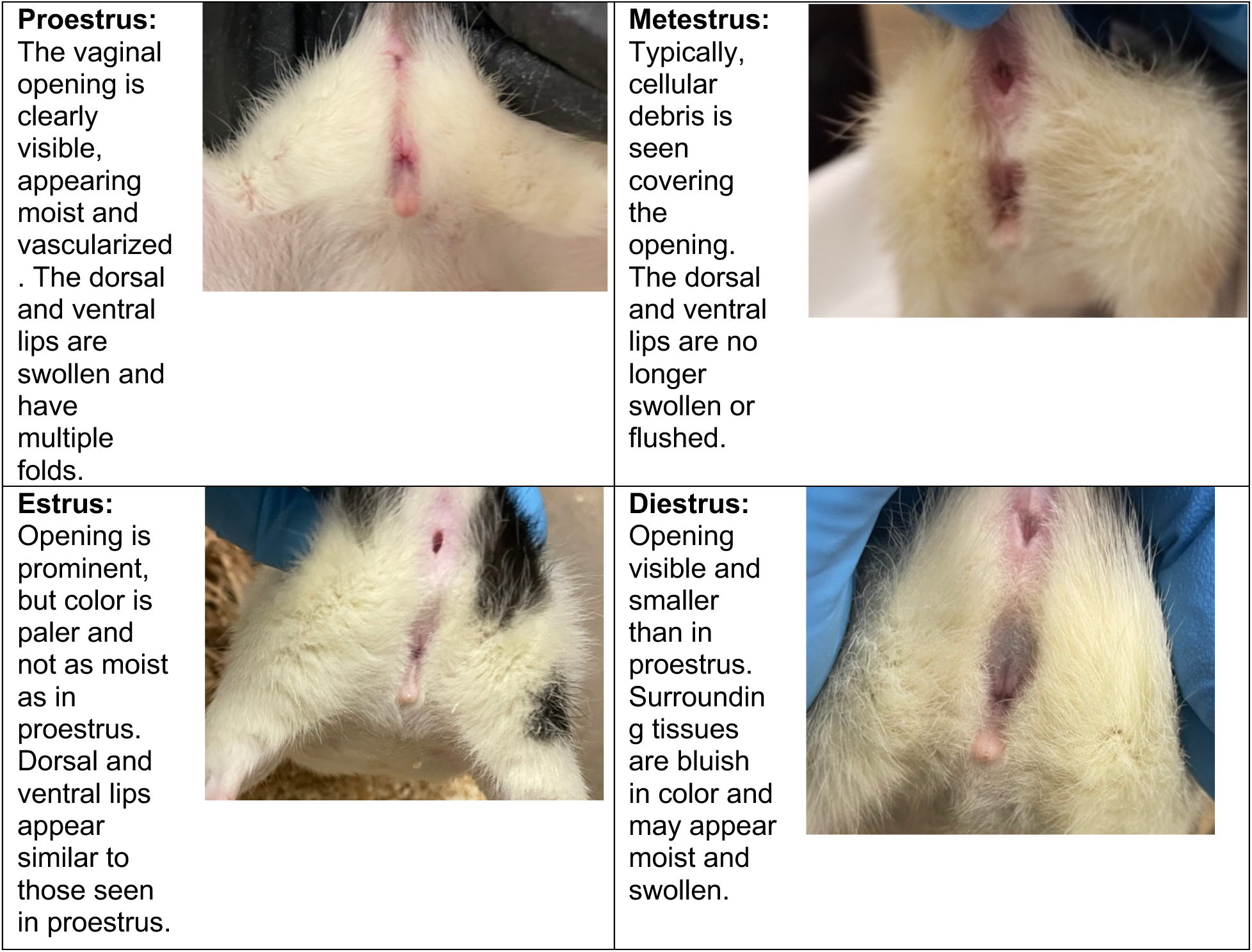
Method for estrus staging.

### Odorants

The monomolecular odorants were methyl salicylate (MS) (TCI Chemicals), 2-nonanone (Thermo Fisher Scientific), ethyl 2-methylbutyrate (EMB; Sigma-Aldrich), 2-hexanone (Thermo Fisher Scientific), and anisole (Fluka). For female and male urine, soiled cage bedding was used. The soiled bedding was collected from cages of rats to which the subjects had no prior exposure and was stored in airtight containers at 4°C. Before each session, odor swabs were placed in these containers for 30 minutes at room temperature after which the swabs were used for odor presentation.

### Data Analysis

#### Sampling Duration Analyses and Statistics

The piezo taps were used to determine sampling duration. Only the data from the first paradigm was used for these analyses because of the difference in protocol for duration of sampling in the second paradigm. Because rats habituate to odors after 2-3 trials, and the first trial was limited to 10s, we used the second trial for sampling duration analysis using a repeated measures 3-way ANOVA with sex, odor, and day as factors.

#### LFP Analyses and Statistics

Signals from all electrode leads for each rat were visually inspected for 60Hz line noise and movement artifact, and the lead with the cleanest signal was used for each rat. Of note, no systematic pattern in the location of good leads was observed across the sexes. A 2-second period preceding the trial onset, henceforth referred to as the pretrial period, was chosen as the trial baseline. Trials with obvious movement artifact were discarded, and only trials with sampling times of at least three seconds were used in electrophysiological analyses. The data from the first second after the trial onset was skipped to eliminate non-uniform odor exposure (the time between the piezo tap and the swab-nose contact was around 1 sec). The next two seconds were taken as the odor trial period (Fig. 2A). Of note, this period is consistent across rats tested in both odor presentation paradigms. For each odor block, we calculated the mean power across trial and pretrial periods. The beta (15-30 Hz) and gamma (65-120 Hz) power ratios for each block were calculated by dividing the mean power across the trials in that block by the mean power across the pretrial periods in that block. If movement artifacts or sampling durations of less than three seconds resulted in fewer than four trials for an odorant in a session, data from that odorant was not included in analysis of the session. Power spectral density analysis was performed using multitaper methods based on the fast Fourier transform (Mitra & Bokil, 2008) with 5 tapers and a time-half-bandwidth of 3 tapers on each data segment (Fig. 2B). Thus, for each animal an 8X12 matrix was obtained corresponding to 8 odors (average odor/pretrial gamma or beta ratio and 12 days). Of note, exclusion criteria for the trials resulted in missing values in the matrix which account for 9.17% of the data for females and 21.67% of the data for males. A repeated measures 3-way ANOVA was conducted to investigate the effect of sex, odor, and day on gamma and beta oscillation power ratios. Where main effects were found, post hoc analyses were applied to determine the direction and significance levels of the differences. Analyses were done with MATLAB® R2021b (Mathworks), and the graphs were generated in MATLAB and on GraphPad Prism 9.4.0.

**Figure 2.**
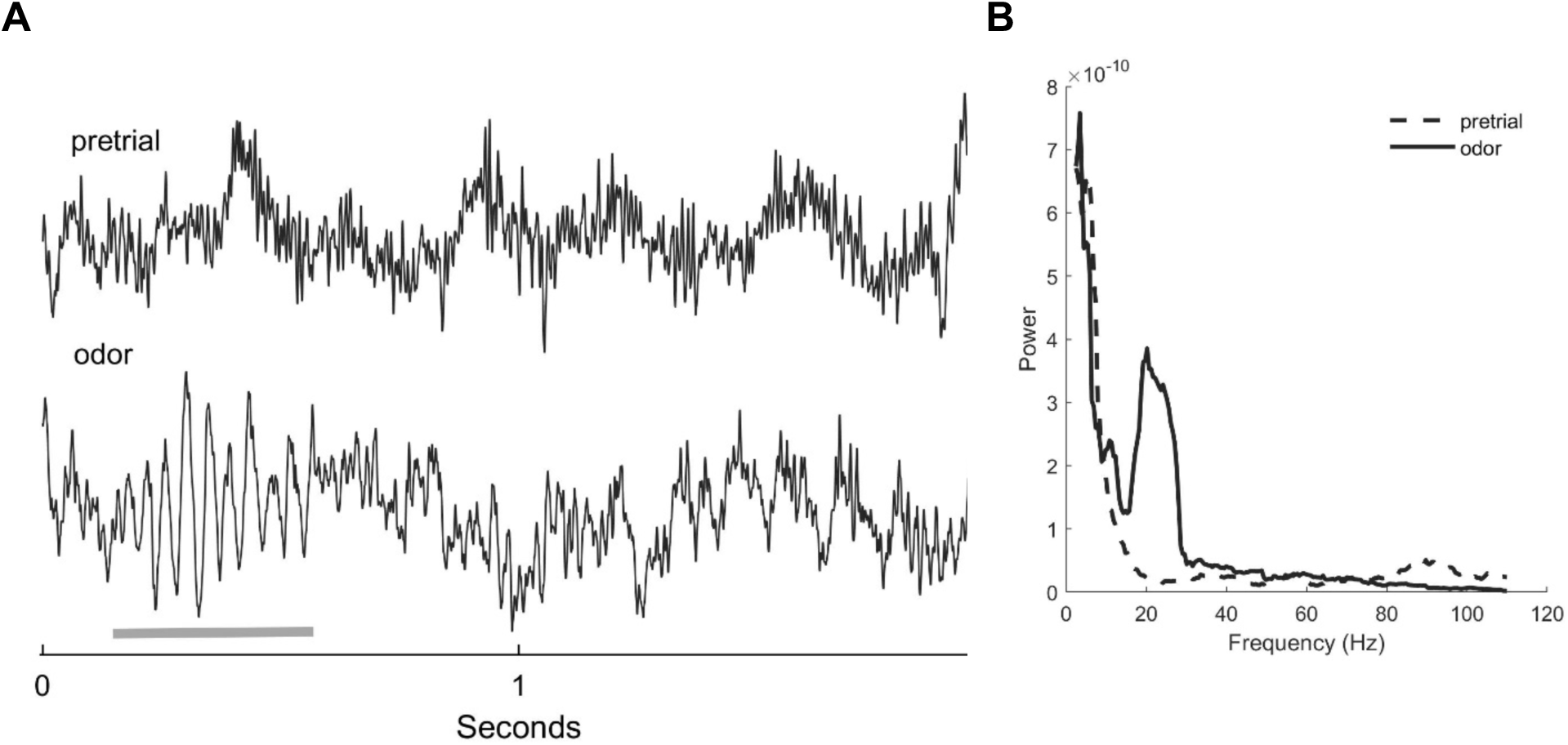
Analyzing oscillation power. **A:** Sample data showing the pretrial period on top and the analyzed section of the odor trial period on the bottom. The X-axis represents time (in seconds) and the y-axis represents voltage (normalized dimensionless signal). Both segments are 2 seconds long and on the same scale. Note the large beta oscillation early in the odor trial where the gray underscore indicates (7^th^ trial on exposure to EMB). **B:** Spectral analysis of the same data seen in left plot, with a prominent beta peak (20-25Hz) and the suppressed gamma (80-85 Hz) during odor presentation. The Y-axis represents power (from normalized data and hence dimensionless with no units) and the X-axis represents frequency (in Hz). Cortical LFP data typically show a 1/f falloff in power, with lower frequencies having logarithmically more power than higher frequencies (Kay, 2014). That pattern is repeated here with a deviation in the quasi-periodic beta band.

#### Baseline Analyses

To check if females and males differ in session baseline oscillatory gamma or beta power, we chose a detrended 6 second period before the start of the experiment each day when the rat was verified to be alert but unmoving. The detrend function in MATLAB computes the mean for the time series data and subtracts that from the data, removing any slow change in the mean power. The average power across beta and gamma band frequency per session was obtained for each rat using the same electrode leads as in the analysis above.

#### Pretrial LFP Analyses

To check if pretrial beta and gamma were different between sexes, we filtered the 2-second pretrial period for obvious movement artifacts, and then pooled detrended pretrial data for each session. The average power across beta and gamma band frequency for each session was obtained for each rat.

#### Estrus Analyses

Estrus staging was performed on each day of recording. From the 12 days of data obtained, two full 4-day estrus cycles were used for analysis, starting with the first observed proestrus that did not occur on day 1 to exclude the effect of the novelty associated with the first day of the experiment. Data before and after two complete cycles were not included for this analysis. A repeated measures 3-way ANOVA was performed on sampling duration and gamma and beta power with sex, odor, and the four days of estrus as factors. Each female rat was paired with a male rat that completed the experiment in the same timeframe and the same analysis was applied to the paired male, using the female’s estrus days to yoke the male’s days for analysis. Thus, an interaction between sex and estrus day, specific to females, would be necessary to indicate an effect of estrus.

## Results

To examine sex differences in odor sampling and olfactory bulb oscillation power, ten rats (5F) were tested across 12 days. In each session (one per day), a rat was exposed in a block design to seven different odors for eight trials each, with a blank trial between blocks. Sampling times on the second presentation of each odor plus averaged beta and gamma power were examined for each of the rats. The beginning of the experiment was not aligned with estrus in the female rats to prevent confounding effects of novelty on estrus day. One female rat was started on day 1 of estrus, one on day 2, two on day 3 and one on day 4.

### Sampling Duration

We used the second trial for each odor to measure sampling time differences. This reduced the effect of odor changes between blocks and still avoided habituation that typically occurs over more trials (see Lowry & Kay, 2007). Six rats were used for this analysis, because the protocol was inadvertently changed for four of the rats (2F). A repeated measures 3-way ANOVA with sex, odor, and day as factors on the second trial for each odor revealed a significant main effect of sex [F(1,384)=24.39; p = 1.17e-6] with a small effect size (η_p_^2^ =0.0373) (Fig. 3). Females (5.799 sec) sampled for significantly shorter durations than males (6.697 sec).

**Figure 3:**
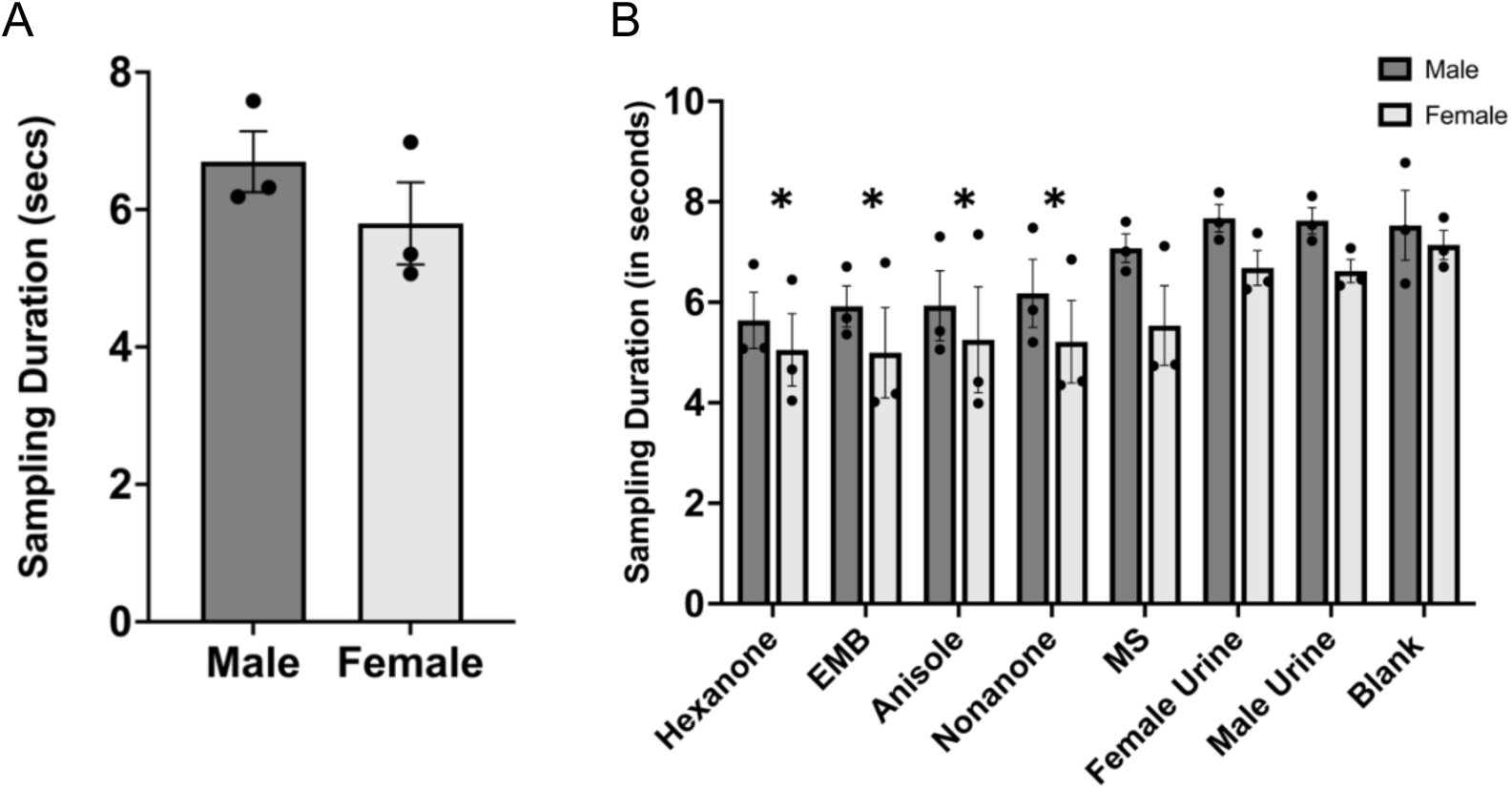
Sampling duration. **A.** Females sampled odors close to 1s less time in trial 2. **B.** Sampling duration across odorants [n=6 (3F)]. The monomolecular odorants are arranged in decreasing order of volatility starting from hexanone and ending with MS. Hexanone, EMB, anisole, and nonanone had significantly lower sampling durations relative to the blank swab. Asterisks indicate a significant difference from blank for the group of 6 rats (no odor by sex interaction). All error bars represent SEM.

The ANOVA also revealed a significant effect of odor [F(7,384)=11.69; p = 1.6*10^-13^] on sampling duration, with a large effect size (η_p_^2^=0.1150). Post hoc analyses (Wilcoxon matched-pairs signed rank test) revealed that hexanone, EMB, anisole, and nonanone were sampled for significantly less time compared to the blank swab. There was no significant sex by odor interaction.

The ANOVA also revealed a significant effect of the experimental day on overall sampling times [F(11,384)=2.64; p = 0.0029], and no significant interactions between day and odor or sex. Post hoc analyses (Fisher’s Protected Least Squares Difference) showed that sampling times dropped significantly on day 11 (lower than days 10 and 7), and the sampling duration for day 12 was significantly lower than that of day 10.

To compare the variances in sampling durations between males and females, we performed an F-test on the median sampling durations averaged across days and odors. The variances between the two groups were not significantly different (F = 1.808; p= 0.7124). Estrus analyses revealed that while the effect of sex (F (1,320) = 18.89; p = 1.85*10^-5^) and odor (F (7,320) = 7.64 p = 1.56*10^-8^) were retained, no effect of estrus day was seen on sampling duration.

### Gamma oscillations

Gamma oscillations represent precision in firing among the population of OB mitral cells that project to other olfactory and limbic brain regions (Rojas-Líbano & Kay, 2008). Increased power in the gamma band increases odor discrimination ability (Beshel et al., 2007; Lepousez & Lledo, 2013; Nusser et al., 2001) A repeated measures 2-way ANOVA on the 6-second session baseline gamma power with sex and day as factors showed no effect of sex (F(1,96)=1.48; p = 0.15) or day of experiment (F(11,96)=0.5; p = 0.65) on gamma power, indicating no significant difference in baseline (not odor-evoked) gamma power between males and females or across days. Pretrial gamma power also did not show differences between sexes or across days. Repeated measures 2-way ANOVA on pretrial gamma power revealed no significant main effect of sex (F(1,96)=2.39; p = 0.13) but a main effect of the day of experiment (F(11,96)=2.24; p = 0.02). (No pairwise day comparisons differed significantly in post hoc analyses. These mostly null effects argue against baseline differences in male and female rats or across days.

Odor-evoked gamma power did show significant effects, falsifying the null hypothesis that male and female OBs do not differ from each other in their oscillatory responses (Fig. 4A,B). A repeated measures 3-way ANOVA with sex, odor (7 odorants plus blank) and day (12) showed a significant small effect of sex (F(1,620)=85.36 p = 3.9*10^-19^; η_p_^2^ =0.0215). Gamma power in females (mean 1.2832, std = 0.6230) was lower than in males (mean 1.8617, std = 1.1788). The ANOVA also revealed significant small main effects of odor [F(7,620)=6.25, p = 4.2*10^-7^; η_p_^2^=0.0028) and day of experiment (F(11,620)=1.88, p = 0.0392; η_p_^2^=0.0009). Post hoc analyses (Wilcoxon matched pairs signed rank test) revealed that hexanone and female urine had significantly lower and higher gamma power respectively, relative to blank. Additionally, post hoc analyses with Fisher’s Protected Least Squares Difference showed that gamma power was not significantly different between any two days.

**Figure 4:**
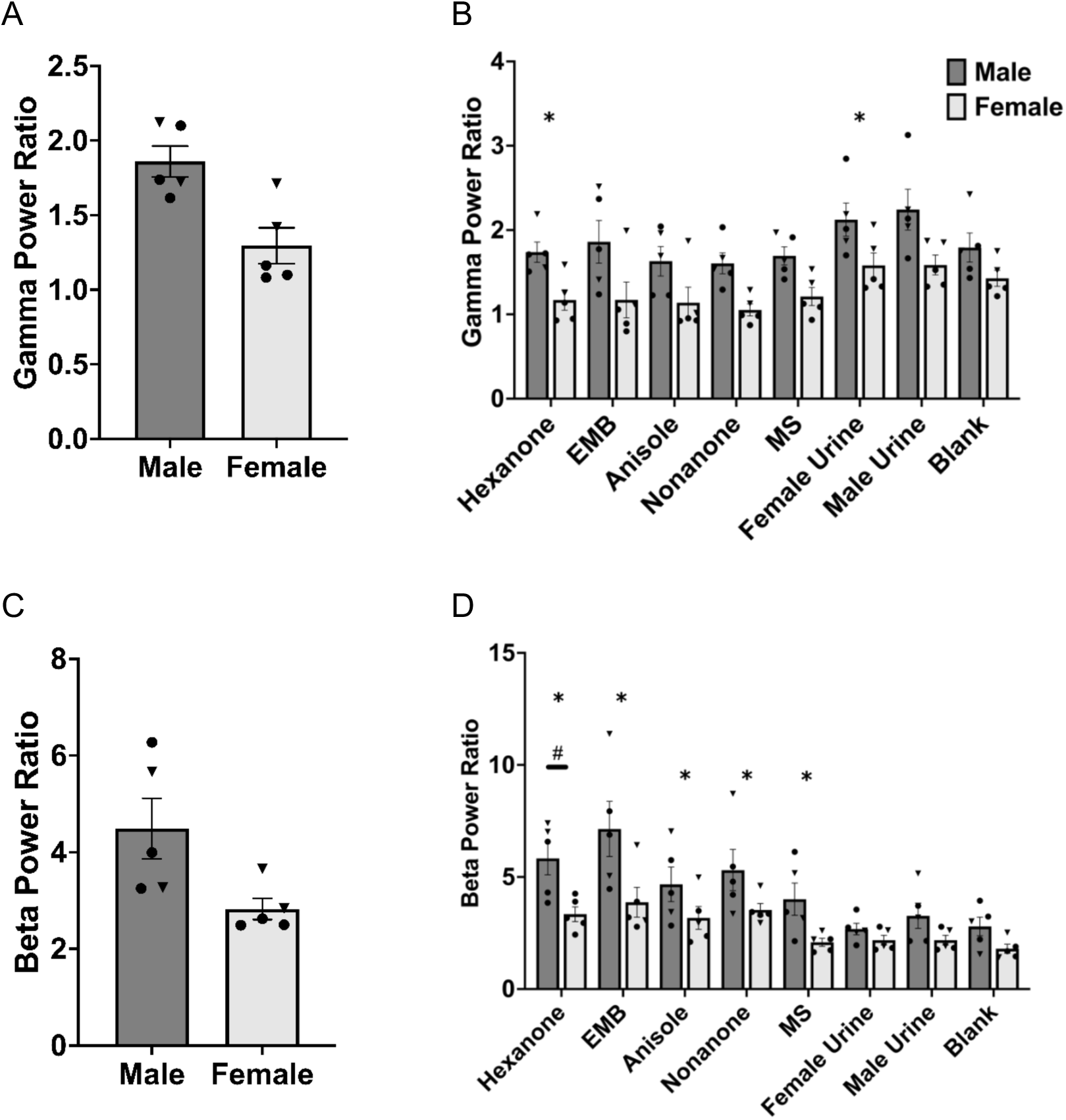
Gamma and beta power effects. **A.** Comparison of gamma power ratios (odor to pretrial) between sexes: Females had significantly lower gamma power ratios than males. **B.** Gamma power ratios of males and females across odorants. Hexanone and female urine had significantly lower and higher gamma power respectively relative to the blank swab (represented by the *). There was no sex by odor interaction. **C**. Comparison of beta power ratio between sexes: Females had significantly lower beta power ratios than males. **D**. Beta power ratios of both sexes across odorants. Hexanone, EMB, anisole, nonanone, and MS had significantly higher beta power ratios relative to the blank swab (represented by the *). The beta power ratio in response to hexanone was significantly higher in males than in females (represented by #). Monomolecular odorants in B and D are arranged in decreasing order of volatility starting from hexanone and ending with methyl salicylate. Urine and blank swabs at the end are on the low end of the volatility scale. Vertical scales for gamma and beta are different due to characteristic decrease in power with increasing frequency (see Fig. 1). Solid circles represent data points from rats in the first paradigm, and the inverted solid triangles represent data points from the rats of the second paradigm.

To compare the variances between gamma power ratios in males and females, we conducted an F-test on the average gamma power ratio for each rat across the 8 odorants and 12 days of experiment. The variances between males and females were not significantly different (F = 1.324; p = 0.79).

Estrus analyses showed that while a significant effect of sex (F(1,486)=61.17; p = 3.3*10^-14^) and odor was retained (F(7,486)= 4.46; p = 8.02*10^-5^), no effect of day of estrus (F(1,486)=1.06; p = 0.36) was seen on odor-elicited gamma power ratios.

### Beta oscillations

Beta oscillations are known to be related to odor volatility in this paradigm, with highly volatile odors inducing larger beta oscillations than low volatility odors (Lowry & Kay, 2007). We tested whether beta oscillations vary in amplitude by sex and whether the volatility effect found previously for male rats was robust across sexes.

We first tested whether there were any baseline differences in beta power. A repeated measures 2-way ANOVA on beta power calculated from the session baseline with sex and day as factors showed no effect of sex (F(1,96)=2.25; p = 0.16) or day of experiment (F(11,96)=0.58; p = 0.50). A repeated measures 2-way ANOVA on the pretrial data with sex and day as factors showed no effect of sex (F(1,96)=3.58; p=0.61) or day (F(11,96)=0.32; p=0.98) on beta power. Both baseline analyses indicate that there are no sex differences in beta outside of odor exposure suggesting that the physiological structure of the OB is likely not changing across days of the experiment.

Odor-elicited beta oscillations did show significant main effects (Fig. 4C,D). A repeated measures 3-way ANOVA with sex, odor (7 odorants plus blank), and day (12) showed a significant small effect of sex (F(1,620)=61.48, p = 1.6*10^-14^; η_p_^2^=0.0148), odor (F(7,620)=15.73; p = 4.6*10^-19^; η_p_^2^=0.0065), and day (F(11,620)=2.51; p = 0.0043; η_p_^2^=0.0011) on odor-elicited beta power. Mean beta power across days and odors was lower for females (mean 2.8072, std = 1.9640) than males (mean 4.4721, std = 4.1375). The ANOVA also revealed a significant interaction between sex and odor (F(7,620)=3.29; p = 0.0019). Post hoc analyses on the beta power ratios of the odors (Wilcoxon matched-pairs signed rank test) indicates that the odors in the high volatility range, hexanone, EMB, anisole, nonanone, and MS had beta power significantly higher than to the blank. Additionally, males showed significantly higher beta power in response to hexanone than females. These results replicate in females our previous findings regarding elevated beta power for high volatility odorants. We also note females showed lower beta power relative to males, similar to the results in gamma power.

Post hoc analyses (Fisher’s Protected Least Squares Difference) showed that the beta power on days 11 and 12 was significantly elevated relative to days 4 and 5 (Fig. 5).

**Figure 5.**
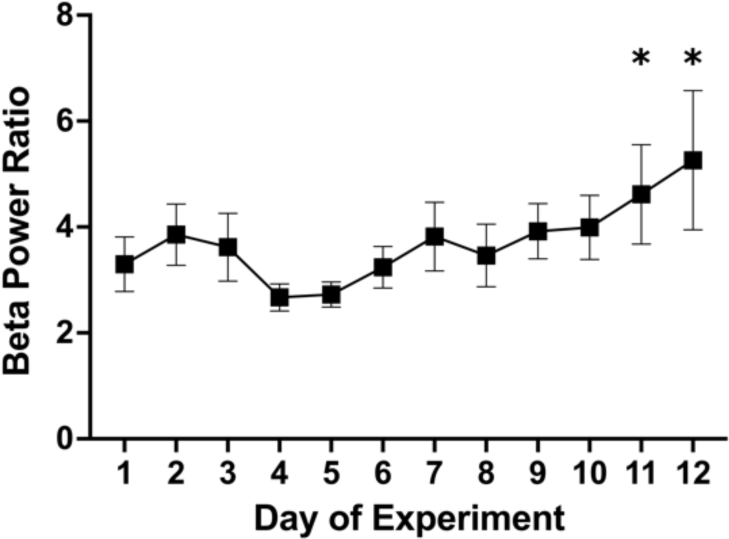
Comparison of beta power ratio across the 12 days of the experiment. Beta power ratios on day 11 and 12 were significantly high relative to days 4 and 5 (represented by the *). All error bars represent SEM.

To compare the variances of beta power in males and females, we conducted an F-test on the average beta power ratio for each rat across the 8 odorants and 12 days of experiment. The variances between the two groups were not significantly different (F = 8.199; p = 0.06).

Estrus analyses revealed an effect of sex (F(1,486)=23.66; p = 1.5*10^-6^) and odor on beta power (F(7,486)=11.68; p = 9.2*10^-14^) and a main effect of day (F(3,486)=3.91; p = 0.0089), but we did not see an interaction between sex and day, suggesting that the estrus day effect is either random or does not influence females differently from males (males and females were not always tested on the same days). We also found a significant interaction between sex and odor (F(7,486)=2.36; p = 0.02) with the beta power ratio for anisole being significantly higher in males relative to females in the estrus analysis.

While gamma power is elevated at the peak of every sniff, except when beta oscillations are present, beta oscillations show a sharp onset and offset (Fig. 2A). Thus, a longer latency to start a beta band oscillatory response to an odor might reduce power within the 2-second window we chose for analysis. To determine whether a latency difference contributed to reduced beta power ratios in females, we checked for differences in the time of beta onset across trials between the sexes. We used a root mean square smoother on the data of the first day of each rat to obtain the earliest point that registered a beta power increase for each trial. The waveforms were inspected and a final threshold of three times the standard deviation of beta power was set to catch beta bursts. Data for day 1 was not available for one of the male rats. A one way ANOVA showed no significant effect of sex on the start time of the first beta burst (F(1,556=1.25 p = 0.26).

## Discussion

Gamma and beta oscillations have been linked to rats’ ability to discriminate, recognize, and make appropriate responses to odor stimuli in operant tasks (Beshel et al., 2007; Frederick et al., 2016; Lowry & Kay, 2007; Martin et al., 2004, 2007; Martin & Ravel, 2014). Beta oscillations have also been linked to odorant volatility in a passive odor exposure paradigm in waking, but not anesthetized, male rats (Lowry & Kay, 2007). We therefore examined the power of gamma and beta oscillations to address the possibility of sex differences in OB neurophysiology which may impact odor perception differences. To our knowledge, ours is the first study to address this question from the perspective of gamma and beta oscillations in the OB. We used a passive odor presentation paradigm to investigate odor sampling time and OB beta and gamma power. We report significant differences between the sexes in sampling behavior and odor-elicited gamma and beta power. The main findings indicate that females sample odors for a shorter time early in an exposure series and, over all twelve sessions, females produce lower power odor-evoked gamma and beta oscillations than males. Importantly, we found no significant differences in beta or gamma power outside of odor sampling during the baseline period at the beginning of the session, suggesting that the sex difference is specific to the role these oscillations play in odor responses. None of the sex differences are attributable to specific estrus days, and male and female rats show equal variance of the measures we examined.

### Behavior

Our study shows evidence for sex differences in odor sampling behavior. We observed that sampling duration in females was shorter than that in males. This aligns with studies in mice showing enhanced spatiotemporal sensitivity of OSNs in females (Kass et al., 2017). The shorter sampling durations in females that we show here (0.89 seconds shorter than males) could be a result of stronger responses to odorants in olfactory receptor neurons, allowing females to sample a comparable amount of odor in less time. We have reported previously that sampling duration showed an inverse linear relationship to the log of the theoretical vapor pressure of the odorants, meaning that stronger odors are sampled for less time (Lowry & Kay, 2007). Consistent with the previous results, we show here that both male and female rats sample only the highly volatile odorants for shorter durations relative to the blank swab. Previous studies on the presence of volatile compounds in urine and differences in attraction thresholds between sexes for urines of both sexes point towards enhanced sensitivity in females (Baum & Keverne, 2002). We did not see a significant interaction between sex and sampling times for the male and female urine odors in our study. Additionally, the sampling durations of both urine swabs were comparable to that of the blank swab. This could be because we used cage bedding as representations of the urine odors. The cages were not on a ventilated rack, but the animal room itself is well-ventilated, and cage changes occur twice-weekly, thereby likely reducing the concentration of volatile odors that would be present in urine. Swabs concentrated with urine collected from rats could potentially elicit stronger behavioral responses. Sampling duration across all odors dropped significantly on days 11 and 12 likely because of long term habituation and loss of novelty of the odors.

### Olfactory bulb oscillations

Olfactory bulb oscillations are some of the most well-studied cortical oscillations, with detailed physiological and mathematical analysis reaching back to Adrian’s papers (Adrian, 1942, 1950). Diverse labs have used either male or female rats and mice and sometimes both, noting similar types of oscillations in both sexes. We have not found any previous comparison of the sexes concerning oscillatory characteristics of OB oscillations. Gamma oscillations are primarily implicated in odor discrimination (Beshel et al., 2007; Stopfer et al., 1997). Better discrimination of similar odorants (fine odor discrimination) corresponds to increased gamma (Nusser et al., 2001), and blocking gamma or the analogous insect antennal lobe oscillation impairs fine odor discrimination (Lepousez et al., 2010; Lepousez & Lledo, 2013; Stopfer et al., 1997). Considering that female rats and humans are known to have better odor discrimination than males (Pietras & Moulton, 1974; Sorokowski et al., 2019), one hypothesis could be that gamma power in females would be expected to be higher. Lower gamma power in females in our study is an interesting observation but not entirely contradictory to what is known about the functional significance of OB gamma. OB gamma power has been shown to increase during discrimination of fine odors and significantly more than during coarse odor discrimination (Beshel et al., 2007). Gamma is produced in the OB within local circuits, and neuromodulator levels likely modulate gamma based on task demands (Chelminski et al., 2017; Osinski & Kay, 2016). It is possible that females’ lower odor thresholds make odor identifications easier without the need to amplify gamma. Our experiment did not challenge odor processing and so may not exhibit the full range of sex differences. Focusing on sex in task-specific paradigms would complement the results from our study to better understand the connection between physiology and differences in olfactory abilities between the sexes.

Females also showed lower odor-elicited beta power than males over the entire set of odors. While post hoc tests showed a sex difference in beta power for one odor (hexanone, with beta power lower in females), the pattern of beta power across odors was similar for males and females. Beta power in response to the two urine odors and the blank swab were significantly lower than in response to the monomolecular odors for both sexes, indicating the absence of high concentrations of highly volatile molecules in these samples. The monomolecular odors in our study that are on the higher end of the volatility spectrum elicited higher beta power. Both of these observations align with prior work from the lab on male rats (Lowry & Kay, 2007), linking high amplitude beta with high odorant volatility for pure odorants. OB beta is implicated in associative odor learning (Kay & Beshel, 2010; Losacco et al., 2020; Martin et al., 2004) and odor-guided motor behavior and odor responses (Frederick et al., 2016; Hermer-Vazquez et al., 2007). Beta may also serve as a tuning signal prior to odor sampling during pre-stimulus periods when a rat shows ‘learned anticipation’ (Kay & Freeman, 1998). We observed that during the course of the experiments, the rats appeared to habituate to the paradigm and showed signs of learning or anticipating odors (behavior not quantified). While we did not see any significant change in gamma power over the 12 days, beta power increased starting on day 4, with days 11 and 12 having significantly higher beta power relative to days 4 and 5.

Because we used the change in gamma and beta power relative to the pretrial baseline (the power ratio) as our measure in these analyses, it was possible that the smaller increase of the gamma and beta power ratio (odor period / pretrial period) in females could be driven by larger baseline gamma. We tested for this and found no differences in either pre-session baseline or pretrial oscillatory power in either band. This further supports a sex difference specific to the odor-evoked physiological response.

What could be the source of such a difference in oscillatory power? Differences in size of the female olfactory bulb could simply mean that a larger population of cells would produce more desynchrony, resulting in a less periodic and therefore lower power signal during odor-evoked oscillations, but this argument would predict that power differences should also occur during baseline periods, which we did not observe. Alternatively, the sex differences we observe could be a result of organizational effects of sex hormones on the developing OB leading to less coordination of mitral and tufted cell spike timing that does not show up until odors are present and attended to. Because we did not find significant variation that depended on estrus, it is unlikely that circulating sex hormones drive the differences we observed. Our results might also be related to sex differences that have been noted in other mammalian species with respect to connections to higher order areas, glomerular and cellular organization, and sensory input (Kass et al., 2017; Waters et al., 2005)

### Sex distributions, variability, and estrus

Recent efforts to improve sex distributions of subjects in biomedical research mean that many more studies use both male and female animals but usually without an explicit design to test sex differences. Our study controlled for some other sources of expected variance to isolate as much as practical differences that might be due to sex. We decreased any variance from circadian effects by running all subjects within a 3-hour window every day. By pairing male and female rats in cohorts we avoided unbalanced effects of any changes in experimental technique, personnel, or protocol through the course of the experiment.

A prior study on mice reported estrus-linked changes in olfactory ability as well as evoked potentials in the OB (Schmidt & Schmidt, 1980). We saw a significant effect of day of testing in sampling duration. To ensure that the effect of the day of experiment and the day of estrus did not confound each other, we included data spanning two estrus-cycles, none beginning on day 1 of the experiment. In our paired analyses, the female rats did not show any estrus-locked behavioral or neurophysiological changes that were different from the males, nor were they any more variable than the male rats in the study. None of our measures correlated with female estrus day over two complete cycles. However, given that there are known differences in olfactory ability that depend on circulating sex hormones (Kass et al., 2017), a longer study period might be required for cyclicity to become discernable. Also, estrus-linked behavioral and LFP changes might emerge when the rat is faced with specific ethological or cognitive demands.

Previous studies have addressed variability in a large number of physiological measures with the result that variance in male and female mice and rats is comparable (Becker et al., 2016; Prendergast et al., 2014). Despite these findings, some researchers still express concern that female variability in circulating sex hormone levels across the days of estrus might lead to more variability in results from female animals. Our study adds olfactory sampling and OB neural oscillations to the list of measures that show equal variance across sexes. If an effect of estrus day is present but not pertinent to the research question, study design can take advantage of the equal variance in males and females by testing subjects on multiple days or randomly with regard to the estrus cycle. Because female rats housed in the same room do not synchronize estrus (Schank, 2001), testing rats on the same day should effectively randomize estrus day. Testing female rats using any of these estrus randomization schemes should not produce results than are any more variable than those from the unknown, possibly periodic, sources of similar variance found in males.

Finally, the sex difference reported here is but a first step in understanding the neurophysiological differences between male and female olfactory bulb function. Future studies should interrogate the cellular and anatomical differences that may support the physiological and behavioral sex differences we report here. In addition, the small but significant effects we report may be further enhanced (or reduced) under conditions of cognitive challenge in odor discrimination tasks.

## Acknowledgments

We thank Huibo Li and Rui He for advice on and assistance with data analysis. This work was supported by an Institute for Mind and Biology Seed Grant to LMK.

